# Genome composition predicts physiological responses to temperature in polyploid salamanders

**DOI:** 10.1101/2024.09.06.611688

**Authors:** Isabella J. Burger, Michael Itgen, Lynn Tan, Parker Woodward, Linet Rivas-Moreno, Tamyra Hunt, Hailey R. Ready, Xochitl G. Martin Geronimo, Robert D. Denton, Eric A. Riddell

## Abstract

Multi-trait analyses can be used to measure the differential performance of phenotypic traits in species complexes. Hybridization within these complexes can result in a mismatch between mitochondrial and nuclear DNA that may lead to reduced performance and acclimation capacity in hybrids. To test the effect of this mismatch on physiology, we compared physiological performance and acclimation capacity of metabolic rate (*V*CO_2_) and total resistance to water loss (*r_T_)* between two sexual *Ambystoma* species and a closely related unisexual lineage. We also separated unisexuals by their unique biotypes to determine how physiology varies with subgenomic composition. We found that unisexual biotypes exhibited phenotypes more like their related sexual species than to other unisexuals. We also found a trade-off between *r_T_* and *V*CO_2_, with increasing *r_T_* resulting in a decrease in *V*CO_2_. Although we did not find evidence for mitonuclear mismatch, our results indicate that genomic composition of hybrids may be a suitable predictor of hybrid trait performance. Multi-trait analyses are imperative for understanding variation in phenotypic diversity, providing insight into how this diversity affects species responses to environmental change.

## Introduction

The formation of hybrids plays a key role in the process of speciation and extinction (Arnold, 1992). Hybrids can positively or negatively impact parental species through gene flow, competition, or the reinforcement of geographic ranges (Arnold, 1992; Barton, 2001; Coyne and Orr, 2004; Edelman et al., 2019). The outcome of these interactions can depend on differential fitness driven by phenotypic variation between hybrids and parental species (Barton, 2001; Thompson et al., 2021; Moran et al., 2021). Hybrids can outcompete one or both parental species by exhibiting beneficial phenotypes (i.e. heterosis; Schluter, 2000; Birchler et al., 2006; Hessenauer et al., 2020). Alternatively, hybrids can experience disadvantages due to Dobzhansky-Muller genetic incompatibilities (DMIs), in which the interactions between divergent alleles can result in reduced performance and, in some cases, hybrid lethality (Dobzhansky, 1982; Arnold and Martin, 2010; Thompson et al., 2023; Moran et al., 2024). Thus, the success of a hybrid often depends upon their performance relative to the parental lineages (Barton, 2001; Thompson et al., 2021). Understanding the effects of hybridization on performance can provide insight into the mechanisms that facilitate or impede hybrid success in species complexes.

The process of allopolyploidization—hybridization events that result in offspring with more than two sets of chromosomes—can reduce performance through the introduction of genetic incompatibilities and fundamental effects on cell size (Gregory and Mable, 2005; Balao et al., 2011; Mable, 2013; Wertheim et al., 2013; Moran et al., 2024). For instance, several key proteins involved in oxidative phosphorylation are comprised of interacting subunits originating from the nuclear and mitochondrial genomes. Thus, divergent nuclear and mitochondrial genomes may result in intergenomic incompatibilities between nuclear- and mitochondrial-encoded protein subunits (‘mitonuclear mismatch’), resulting in a reduction in mitochondrial efficiency and capacity (Harrison and Burton 2006, Toews and Brelsford, 2012; Wolff et al. 2014), which in turn can lead to decreased performance (Sackton et al. 2003; Comai, 2005; Lee-Yaw et al. 2014; Denton et al. 2017; Rank et al. 2020; Klabacka et al., 2022; Moran et al., 2024) and fitness in hybrid offspring (Yee et al. 2013; Wolff et al. 2014). Therefore, mitonuclear mismatch influences hybrid success by shaping physiological performance.

A secondary consequence of allopolyploidization is its effects on DNA content and, consequently, cell size (Gregory, 2001; Balao et al., 2011). The total amount of DNA within a cell (i.e., genome size) is strongly correlated with the size of the cell; therefore, polyploidization causes a universal increase in cell size throughout the organism (Hanken and Wake, 1993; Gregory, 2001; Itgen et al., 2022). Large genome and cell size have been hypothesized to drive the evolution of physiological traits through the scaling effects of lower surface-area-to-volume ratios in larger cells (Szarski, 1983; Vinogradov 1995; Kozłowski et al., 2003). Although broad-scale taxonomic studies have found conflicting evidence for this relationship (Licht and Lowcock, 1991; Uyeda et al., 2017; Gardner et al., 2020), there is empirical support for a cell size-mediated role in determining metabolic rate. For example, larger cell size via polyploidy has been shown to lower the metabolic cost of cells and, by extension, larvae in polyploid *Xenopus* frogs by reducing the energetic costs of maintaining ion gradients (Cadart et al., 2023). In addition, organisms with larger cell sizes may have thicker skin, which reduces desiccation risk by lowering water loss rates (Feder and Burggren, 1985; Johnson et al., 2021). Therefore, allopolyploidization might introduce two key consequences (i.e., cytonuclear incompatibility and cell size) that affect physiological performance. Investigating the effects of allopolyploidization on physiological traits is particularly important because these traits are highly integrated and shape variation in suites of behavioral and life history traits that influence performance and fitness (Ricklefs and Wikelski 2002; Wikelski et al. 2003; Martin et al. 2006).

Metabolic rate and water loss are two important physiological traits that covary in terrestrial animals (Woods and Smith, 2010). Specifically, these traits are linked by the properties of respiratory surfaces (e.g., lungs and skin) that facilitate the diffusion of gases, like oxygen and carbon dioxide. Respiratory surfaces are thin and highly vascularized, which enables oxygen to diffuse rapidly across the tissue (Czopek, 1962; Feder and Burggren, 1985). However, the properties that facilitate oxygen uptake also result in high rates of water loss (Feder and Burggren, 1985). This relationship between gas exchange and water loss results in a trade-off: physiological changes to reduce water loss can also limit an organism’s ability to breathe (Addo-Bediako et al. 2001; Woods and Smith, 2010; Riddell et al. 2018a). By lowering water loss rates (e.g., as an acclimation response to desiccation), organisms often experience a concurrent reduction in metabolic rate due to lower rates of gas exchange (Anderson, 1970; Riddell et al. 2018a). This reduction in metabolic rate might lower an organism’s ability to fuel energetically demanding activities, such as foraging, reproduction, and territory-defense (Burggren and Vitalis, 2005; Biro and Stamps, 2010; Sadowska et al., 2015; Sokolova, 2021). However, the strength of the trade-off can vary across species and hybrids (Burger et al., 2024), which may allow these two linked traits to evolve more independently in different lineages. The relationship between these two traits can also be evaluated through the water-gas exchange ratio (WGER) – the ratio of oxygen uptake to water lost – that provides an index of an organism’s respiration efficiency (Burger et al., 2024). Understanding these linkages may help to predict organismal and evolutionary responses to environmental change.

In this study, we explore the physiological consequences of allopolyploidization in a lineage of allopolyploid salamanders (genus *Ambystoma*). We measured the thermal sensitivity of metabolic rate, water loss rate, respiration efficiency, and the trade-off between water loss and oxygen uptake in the unisexual *Ambystoma* complex. We also investigated the acclimation capacity (defined as the capacity of an organism to reversibly adjust a physiological trait in response to a new environment) of these traits because acclimation provides insight into a species’ ability to respond to environmental change (Riddell et al. 2018b). Specifically, we compared traits between two sexual, diploid *Ambystoma* species (Jefferson salamander [*A. jeffersonianum*] and blue-spotted salamander [*A. laterale*]) and unisexual, polyploid salamander hybrids (Figure 1). We hypothesize that the unisexual allopolyploid salamanders will differ in physiological performance compared to the sexual species due to the consequences of allopolyploidization (i.e., cytonuclear incompatibility and large cell size) (see *Study species* in Methods). Due to the potential for cytonuclear incompatibilities, we predicted that unisexual salamanders would have lower metabolic rates, respiration efficiency, and acclimation capacity compared to *A. jeffersonianum* and *A. laterale*. Due to larger cell sizes (as a result of increased ploidy), we predicted that unisexuals will exhibit lower metabolic rates as well as water loss rates (Johnson et al., 2021; Cadart et al., 2023). Analyzing multiple facets of an organism’s ecophysiology will allow us to develop a more comprehensive understanding of the effects of hybridization and environmental variation on whole-organism performance.

**Figure 1.**
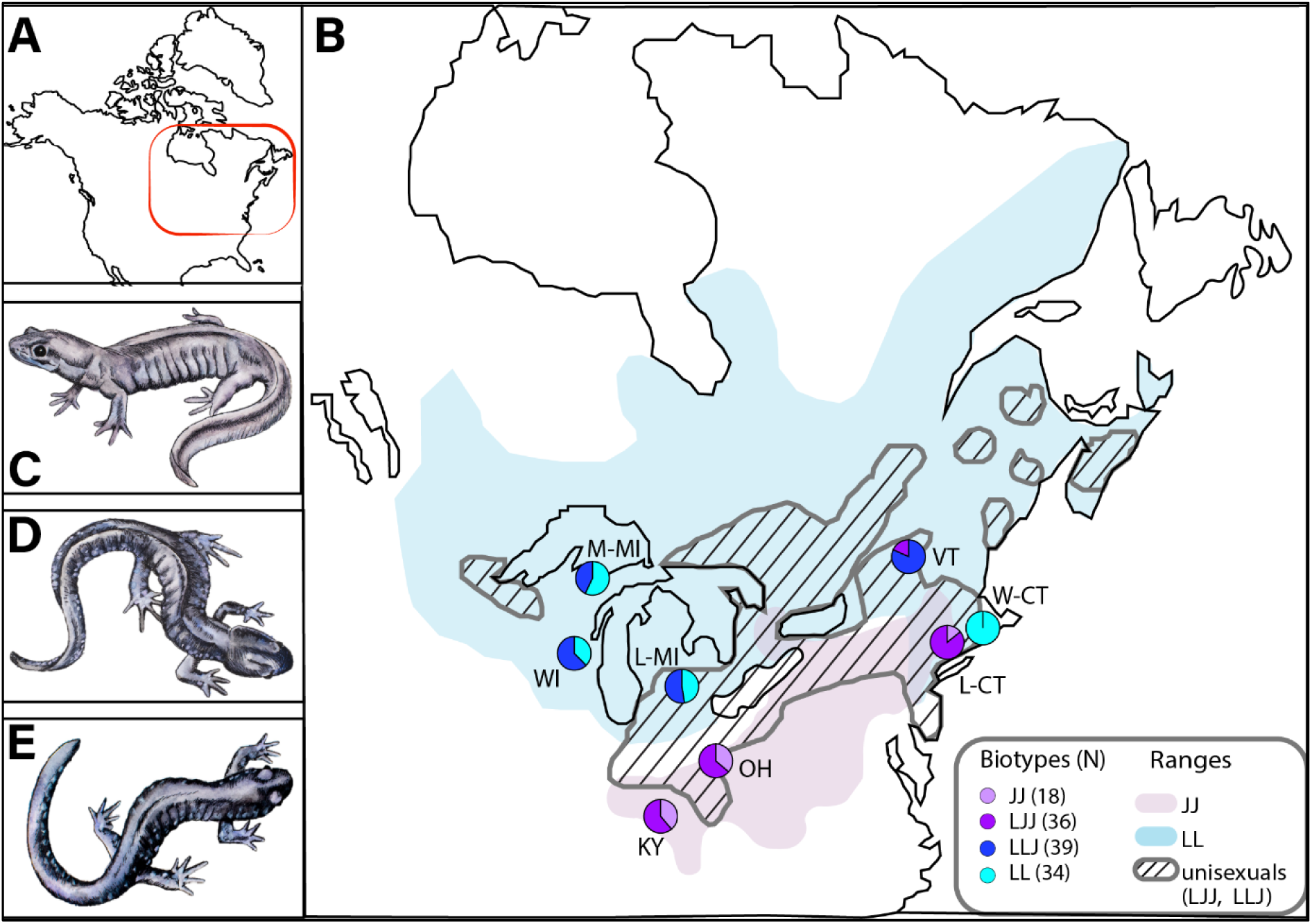
Inset of United States and Canada with the area of interest outlined (A). Area of interest showing distribution, sample size (N), and range of different biotypes across collection sites. M- MI: Marquette, Michigan; L-MI: Livingston, Michigan; WI: Calumet, Wisconsin; KY: Campbell, Kentucky; OH: Delaware, Ohio; VT: Addison, Vermont; W-CT: Windham, Connecticut; L-CT: Litchfield, Connecticut; JJ: *Ambystoma jeffersonianum;* LJJ and LLJ: different *Ambystoma* unisexual biotypes; LL: *Ambystoma laterale.* Pie charts indicate relative numbers of individuals collected at each site (B). Pictures of *Ambystoma jeffersonianum* (C), unisexual polyploid (D), and *Ambystoma laterale* (E). Illustrations provided by Katie Garrett.

## Materials and methods

### Study species

Unisexual *Ambystoma* are an all-female salamander lineage that is the oldest recognized unisexual vertebrate group, persisting for ∼5 million years (Bi and Bogart, 2010). Unisexuals reproduce via kleptogenesis, in which they must “steal” sperm from a congeneric male to stimulate egg development (Bogart et al. 2007; Gibbs and Denton, 2016). The genetic material from males can then either be discarded (offspring are clones of mother) or incorporated into the chromosome array of the embryos. This process has resulted in a range of polyploidy biotypes (2N-5N) that combine genomes from five different species, but the majority of populations are triploid with genomic contributions from *A. laterale* and *A. jeffersonianum* (Robertson et al. 2006; Bi and Bogart, 2010, Bogart et al. 2007). While frequent genome introgression from sexual species into unisexual populations has produced highly variable combinations of nuclear genomes (Bogart, 2019; Gibbs and Denton, 2016), the mitochondrial genome is strictly inherited matrilineally and monophyletic (Hedges et al., 1992; Spolsky et al., 1992). The lack of co-evolutionary history between the mitochondrial and nuclear genomes may result in mitonuclear mismatch and likely has energetic consequences for unisexual salamanders by reducing endurance relative to sympatric sexual species (Denton et al. 2017). Additionally, triploid unisexuals have larger cells than diploid individuals (Licht and Bogart, 1987), and cell size has been linked to variation in metabolic rate and water loss rates in amphibians (Johnson et al. 2021; Cadart et al., 2023; but see Licht and Bogart, 1990). However, studies have yet to compare unisexual biotypes with related sexual species across their range (Figure 1) to make comprehensive comparisons and understand if unisexual biotypes exhibit phenotypes more like the sexual species to which they are most genetically similar. In this study, we compare a suite of physiological traits between *A. jeffersonianum, A. laterale* and unisexuals from the same populations with different genomic composition to determine if mitonuclear mismatch affected unisexual physiological performance. Each of these ‘biotypes’ is described by using the first letter of the parental species epithet, e.g. a LLJ individual is triploid with two sets of chromosomes from *A. laterale* and one set of chromosomes from *A. jeffersonianum*.

### Animal collections and housing

We collected animals between February and April 2020, 2022, and 2023 during breeding migrations. We collected *Ambystoma jeffersonianum* from Litchfield, CT; Campbell, KY; and Delaware, OH; *A. laterale* from Calumet, WI; Marquette, MI; Livingston, MI; Windham, CT; and Addison, VT; and unisexuals from each location (Figure 1B, see figure for sample sizes). These sites were chosen to maximize representation across the range of the unisexual polyploids and target previously surveyed sites that have biotypes most representative of the lineage (triploids, either LJJ or LLJ). We captured sexually mature individuals using a combination of nighttime surveys, terrestrial drift fences, bucket traps, and/or cover boards, depending on the approved collection methods for each state. We acquired scientific collection permits where required (KY:SC2311177, MI:FSCP12302022102023, OH:18-164/SRP23-01, VT:SR-2023-03, WI:SRLN-23-46) and worked with local field personnel when possible to help state agencies as part of larger survey efforts.

We housed salamanders in individual clear boxes (19 x 33 x 10 cm) within refrigeration units set at 12-14°C immediately following collection. Each container contained moist paper towels that were changed weekly and a PVC hide. We fed animals 1-2 appropriately sized European Nightcrawlers (*Eisenia hortensis*) per week. We covered the front glass of the refrigeration units except for narrow slits that would let in ambient light from the room during normal building hours (∼9am-5pm). We weighed each animal once per month and checked animals each day to monitor for any signs of stress. The animal care protocol identified signs of stress as general lethargy, unresponsive to stimulus, weight loss, or unusual skin coloration. None of the animals in this study demonstrated those behaviors, and therefore animals were not removed from our final dataset at this stage (see below for removing salamanders that were active during respirometry trials). Prior to the initiation of this study, we acquired approval from IACUC (protocol #RD-2021-1R). In May 2023, we transported salamanders to [university redacted for double-blind review] for physiological experiments. For transport, we stored individuals in smaller plastic containers lined with dampened paper towels. Total transport lasted approximately 7.5 hours, at which time we immediately removed salamanders from the vehicle and transferred them to a walk-in environmental chamber (2.5 x 3 m) set to 14°C for the duration of the experiment (except for the acclimation trials – see *Physiological experiments*). We kept individuals in the same plastic containers described above. We ensured that lights were kept off in the housing unit, but salamanders were exposed to passive light through a small access window during the day. We made all efforts to keep the housing at the experimental laboratory as similar as possible to that of the location where animals were housed after capture.

### Genetic analysis

We clipped a small tissue sample from the tail of each captured animal to determine their genetic identity. First, we sequenced a highly variable portion of the mitochondrial genome and compared it to known reference sequences for multiple *Ambystoma* species to determine species identity (primers F-THR and R-651; McKnight and Shaffer, 1997; Bogart et al. 2007). For individuals that were identified as unisexuals, we then used fragment length data from three microsatellite loci to determine their specific ploidy and subgenome composition (*following* Teltser and Greenwald, 2015). These subgenomes make up the separate unisexual biotypes (LLJ, LJJ, etc.). One microsatellite locus amplifies only in *A. jeffersonianum*, whereas the other two amplify in both species at different fragment size ranges. As in other studies (Bogart et al. 2007; Bogart et al. 2009; Julian et al. 2003; Ramsden et al. 2005), this approach identifies subgenome composition and can be validated across multiple individuals from the same site due to most polyploid individuals in the same populations sharing the same genomic composition (Bogart and Klemens 2008). Finally, the sites used for sample collection have undergone previous surveys, establishing *a priori* expectations for the biotypes located there.

### Study design

We conducted physiological experiments from May to July 2023. Prior to experimentation, we separated salamanders into five batches of 28 individuals. We further divided these batches into groups of seven, as only seven salamanders could be placed in the flow-through system at a time. We randomized each group by site and species to ensure equal representation in each batch. Salamanders were given one week at 14°C to adjust to laboratory conditions prior to taking physiological measurements in the flow-through system (description below).

We exposed each salamander to three acute temperature treatments in the flow-through system that represent a typical range of spring temperatures they would experience during active periods in the wild: 6°C, 14°C, and 22°C (Lowcock et al., 1991; Hoffman et al., 2018; Mitchel, 2020). To ensure that salamanders reached thermal equilibrium prior to the run, we placed a group in the incubator (Percival, Inc.; Model #I-36VL) at the experimental temperature for one hour prior to physiological measurements. We ran two groups (14 salamanders) a day, and individuals were given a rest day after each run to allow time for rehydration. We determined if a salamander was rehydrated by monitoring their mass prior to and following the respirometry trials. Therefore, it took two days to run a full batch at one temperature.

After the initial physiological measurements at each temperature, we moved the salamanders to a separate incubator (Percival, Inc.; Model #I-36VL) set to 20°C for 20 days to evaluate their capacity to acclimate by adjusting physiological traits. During this time, we increased the rate of feeding to twice per week. We conducted the same physiological measurements following this 20-day period to determine acclimation capacity of each individual. For example, we ran a batch of salamanders at 6□, placed them in the 20□-acclimation chamber for 20 days, then measured their physiology again at 6□ We compared physiological performance between the two 6□ measurements (before and after exposure to the warm acclimation chamber) to determine acclimation capacity. We conducted all physiological measurements between 7:00 and 17:00 to measure physiological traits when salamanders would be inactive in the field (McNab, 1997). Additionally, we stopped feeding salamanders one week prior to experimental treatments to ensure a post-absorptive state. We ensured each individual was at rest during the trials using a Reolink Network Video Recorder (Model RLN8-410) to monitor and record salamander activity. We checked that salamanders were inactive through visual inspection of the data output and verification using the video recordings. Activity is characterized by erratic peaks in metabolic rate. Using this information, we removed all trials where activity was clear and verified inactivity in the other trials using the recorded videos. We removed all trials where the salamanders were active from the final dataset.

### Flow-through system

In preparation for the physiological trials, we placed salamanders on a hardwire mesh platform in a glass chamber (20 x 3.6 cm; volume *c*. 207 mL). Our primary objective was to understand physiological responses to temperature; thus, the mesh restricted the salamander’s ability to behaviorally reduce water loss by curling onto themselves. We placed chambers in a temperature- and humidity-controlled incubator (Percival, Inc.; Model #I-36VL) and connected to a flow-through system (Sable Systems International (SSI), Las Vegas, NV). We delayed physiological measurements by 30 minutes once the chambers were connected to allow the salamanders to come to rest in the new environment.

For the 14°C and 22°C treatments, we used a bench pump (PP-2; SSI) to pull air from the incubator. We also adjusted the relative humidity at each temperature to ensure a similar vapor pressure deficit (VPD) across the treatments. The VPD is the difference between the saturated vapor pressure (the amount of water the air can hold) and actual vapor pressure (the amount of water in the air). As temperature increases, saturation vapor pressure increases, potentially exposing organisms to higher VPDs (and thus greater evaporative demand) (Riddell et al. 2017, Weaver et al. 2022). By maintaining the same VPD across temperature treatments, we ensured that variation in water loss rates could be attributed to organismal responses to temperature and not the effects of humidity. We adjusted humidity across these temperatures to maintain a VPD of 0.75 ± 5% kPa. We were unable to use the incubator to control humidity at 6°C, so we split the airstream being pulled by the pump into two columns: one pulling air straight from the incubator and one pulling air from the incubator and through a column of Drierite^TM^. We adjusted the ratio of these two columns until we reached the target VPD (0.75 kPa).

After air was pumped out of the incubator, it passed through a flow-through manifold (SSI) and separated into different streams with a continuous flow rate of 100 mL·min^-1^. These streams passed through the chambers containing the salamanders, as well as through an additional, empty chamber that we used to collect baseline data. A multiplexer (MUX8; SSI) cycled through the eight chambers (seven experimental and one baseline), and excurrent air samples from each passed through a water-vapor analyzer (RH300; SSI) and a carbon dioxide analyzer (CA-10A; SSI) to measure water vapor and CO_2_, respectively, in volts. We calibrated the analyzers once a month throughout the duration of the experiment to ensure accurate data collection. We calibrated the CA-10 by passing N_2_ gas and a known concentration of CO_2_ gas through the analyzer to zero and span the system, respectively. We calibrated the RH-300 by passing N_2_ gas through the system to set zero and using our humidity-controlled incubator to set the span. The empty chamber was sampled between each salamander measurement to correct for any drift throughout the duration of the experiment. We measured individuals three times at five-minute intervals over a three-and-a-half-hour period.

We connected each instrument to a universal interface (UI-3; SSI) that uploaded the voltage data to Expedata. We then linearly transformed the data in Expedata using equations provided by the manufacturer (SSI). We used lag- and drift-corrections to correct for the time delay in gas exchange/water loss and for drift that occurred during the experiment, respectively. We also corrected the flow rate and carbon dioxide measurements for barometric pressure and water vapor pressure of the excurrent air (Lighton 2008). We then transformed CO_2_ values from percentages into partial pressure by dividing the corrected data by 100. We measured the asymptote of water loss values and used z-transformations to calculate stable water loss measurements. We used equations 3-5 in Riddell et al. (2017) and 10.4 in Lighton (2008) to calculate evaporative water loss (EWL), total resistance to water loss (*r_T_*), and metabolic rate (*V*CO_2_) using the collected data. We measured total resistance to water loss as the inverse of water loss, while accounting for the evaporative demand of the air (Riddell et al. 2017). We averaged *V*CO_2_, *r_T_*, and EWL values across the measurements when salamanders were at rest.

We evaluated the relationship between gas exchange and water loss using two metrics: the water-gas exchange ratio (WGER; *V*CO_2_[mol]:EWL [mol]) and the trade-off between oxygen uptake and resistance to water loss. The WGER is calculated as the rate of oxygen consumption per unit water loss, providing an index of an organism’s respiration efficiency from a water loss perspective (Burger et al., 2024). The trade-off between oxygen consumption and resistance to water loss more effectively captures physiological changes to minimize water loss (e.g., vasoconstriction of the skin, breathing rate; Riddell et al., 2019) and how these changes affect oxygen uptake. However, resistance controls for the evaporative demand of the air and therefore does not reflect the total amount of water loss; thus, we propose that analyzing both WGER and the trade-off between oxygen uptake and resistance provides the most comprehensive understanding of the relationship between gas exchange and water loss (Burger et al. 2024).

### Statistical analysis

We ran three groups of analyses on the collected data: analysis of the traits, change in the traits (acclimation), and the trade-off between resistance to water loss and gas exchange. For the trait analyses, we ran three separate linear mixed effects models (LMER) to determine how *V*CO_2_, *r_T_*, and respiration efficiency (WGER) varied throughout the experiment (*A. laterale*: n = 34; *A. jeffersonianum*: n = 18; unisexual: n = 80). We included mass as a covariate and species, temperature treatment, and time period (before and after acclimation) as factors in each separate analysis. We also included interactions between species, temperature treatment, and time period to assess differences between species. We included time period as an effect in these analyses to account for the variation introduced by this treatment; however, we restricted our discussion of the effects of acclimation to the change in trait analyses (see below). *V*CO_2_ and WGER values were log-transformed to meet assumptions of normality. For the change in the traits, we conducted additional analyses that used the change in *V*CO_2_, *r_T_*, and respiration efficiency between acclimation periods as the response variables (Δ*V*CO_2_ [*A. laterale*: n = 32; *A. jeffersonianum*: n = 18; unisexual: n = 76], Δ*r_T_* [*A. laterale*: n = 32; *A. jeffersonianum*: n = 18; unisexual: n = 75], and ΔWGER [*A. laterale*: n = 32; *A. jeffersonianum*: n = 18; unisexual: n = 75], respectively). Acclimation was calculated as the change in trait value after exposure to the 20□-acclimation treatment minus the trait value before exposure to the acclimation treatment for each individual at each temperature treatment. For these analyses, we only included data from individuals that had a value for both pre- and post-acclimation. We also removed time period as a factor since we calculated the change in traits across the time periods. We conducted these two groups of analyses (trait and change in the trait) to explicitly test our hypotheses on trait performance and acclimation capacity. Additionally, we used a separate linear regression to evaluate the trade-off between the change in *V*CO_2_ and the change in *r_T_* [*A. laterale*: n = 32; *A. jeffersonianum*: n = 18; unisexual: n = 75]. All models included mass as a covariate and salamander ID nested inside batch number and year collected as random effects. Year collected was removed from models when it did not account for any additional variation in the data. We were unable to include collection site in the above models due to its confounding relationship with biotype (specific biotypes are only found in specific sites, Fig. 1); thus, we ran additional analyses for each individual biotype to evaluate the effect of collection site on trait performance (Appendix 1) and acknowledge that collection site may influence physiological rates in our global models.

We were also interested in whether *V*CO_2_, *r_T_*, and respiration efficiency differed between the unisexual biotypes. Therefore, we ran additional analyses on the traits above but included biotype as a fixed effect as opposed to species (analysis of traits: LLJ: n = 39; LJJ: n = 36; trade-offs: LLJ: n = 37; LJJ: n = 34). In these models, we treated biotype (*A. jeffersonianum* [JJ]*, A. laterale* [LL], LJJ, and LLJ), temperature, and time period as factors. For the change in the trait analyses (Δ*V*CO_2_ = LLJ: n = 37; LJJ: n = 35; Δ*r_T_* = LLJ: n = 37; LJJ: n = 34; ΔWGER = LLJ: n = 37; LJJ: n = 34), we removed time period as a factor (similar to the above analysis). We also included mass as a covariate and the same random effects as the models described above. Sample size of *A. jeffersonianum* and *A. laterale* were the same as above for all biotype analyses. If any analysis was significant, we conducted pairwise analyses using the emmeans package to investigate the differences in traits between specific biotypes. We then compared the species-based models with the biotype-based models using AIC to determine which set of models were a better predictor for physiological performance.

Additionally, we calculated ω^2^ to measure the effect size of each variable in each analysis using the following equation:

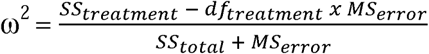

where *SS_treatment_* is the sum of squares for the parameter, *df_treatment_* is the degrees of freedom for the parameter, *MS_error_* is the mean square error, and *SS_total_* is the total sum of squares (Olejnik and Algina, 2003). We also calculated marginal and conditional *R*^2^ to describe the variation accounted for by the fixed effects and the variation accounted for by all effects in the model (fixed and random), respectively. We show all data as mean ± SEM using the *effects* package in R, which returns an adjusted mean incorporating the effects of all variables in the model. We conducted all analyses in R (version 4.3.1). Table 1 reports the outputs from the above linear mixed effects models, and Table S1 reports the outputs from the pairwise comparisons in our biotype models.

**Table 1:**
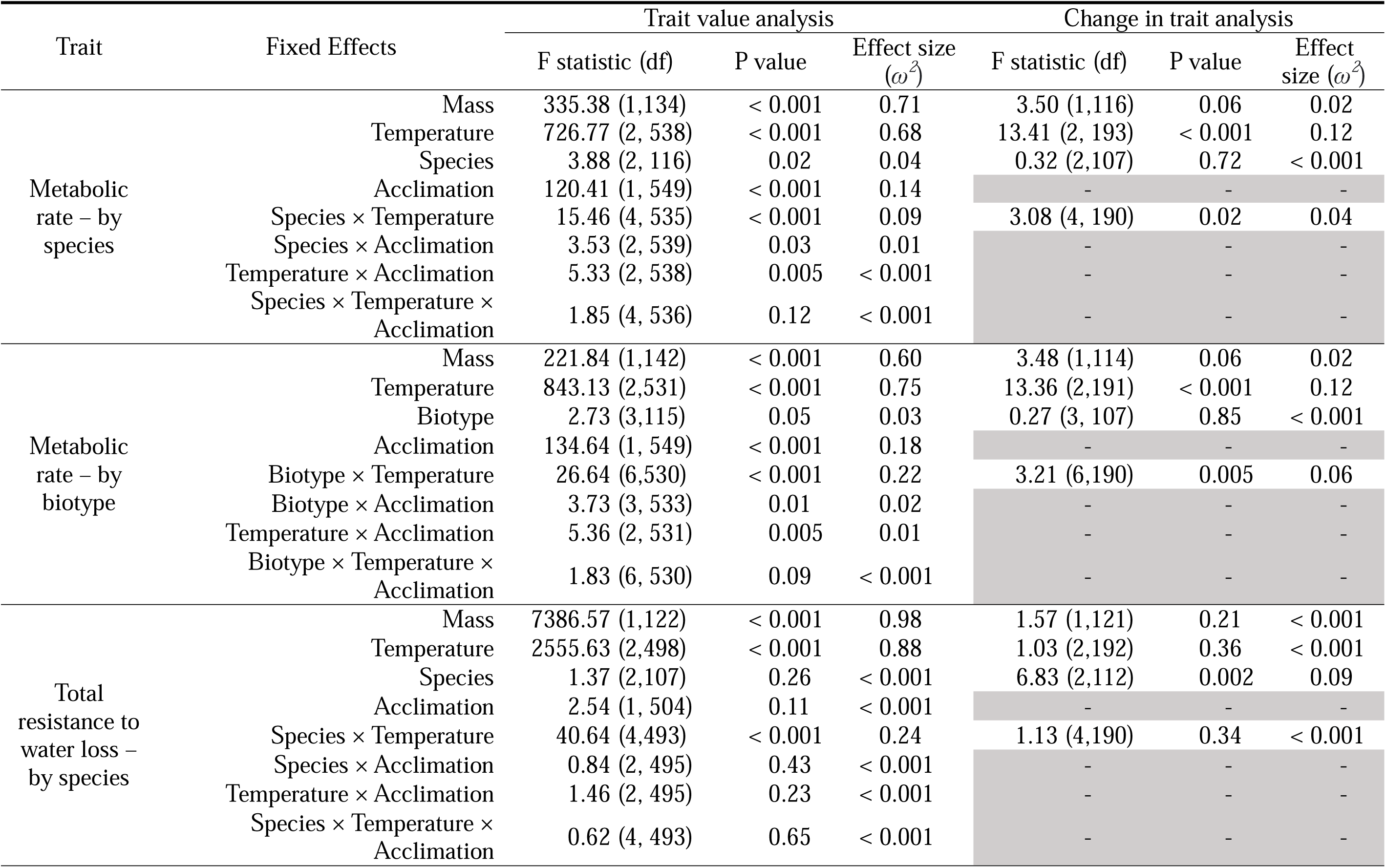
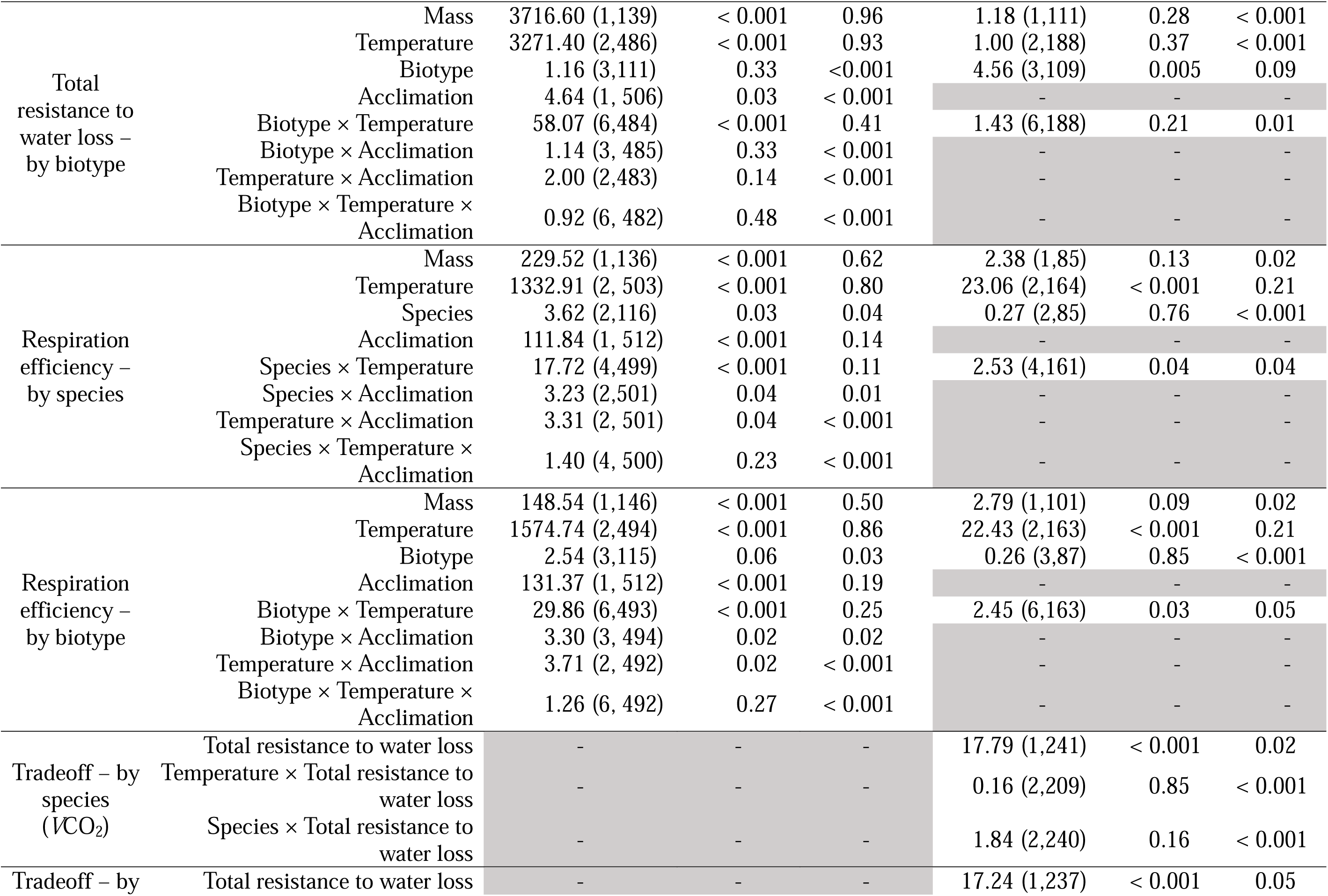
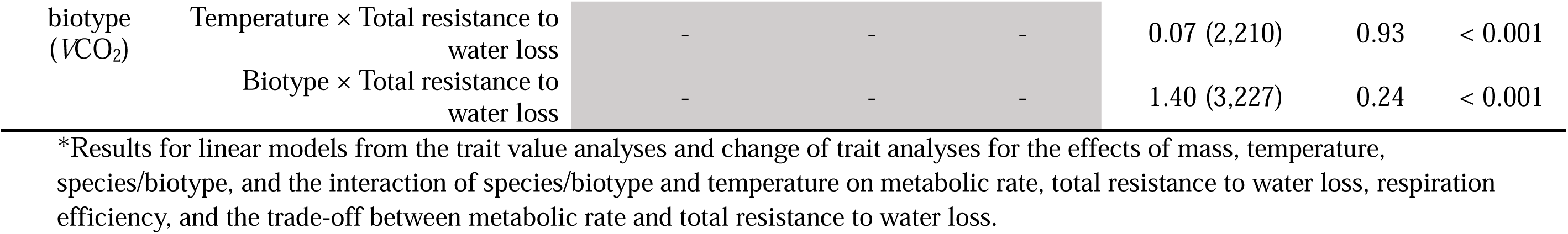
Results for linear models from the trait value analyses, change of trait analyses, and trade-off analysis*.

## Results

### Metabolic rate (VCO_2_)

Our results indicated that *V*CO_2_ was positively related to mass, with larger individuals having higher metabolic rates (see Table 1 for all statistics). We also found that *V*CO_2_ increased with temperature, and *A. jeffersonianum* had a higher *V*CO_2_ than *A. laterale* and unisexuals. The interaction between temperature and species also influenced *V*CO_2_, with *A. jeffersonianum* having the highest sensitivity to the changes in temperature (Figure S1A). Moreover, 78% of the variance was explained by our treatments and mass as indicated by the marginal *R*^2^. In the Δ*V*CO_2_ analysis, salamanders reduced metabolic rate by 19% following the 20□-acclimation period (Figure S1B). Additionally, Δ*V*CO_2_ was positively related to mass and was greatest at 22°C. There was also an effect of the interaction between temperature and species on Δ*V*CO_2_, with *Ambystoma jeffersonianum* having the lowest change in metabolic rate at 6°C but the highest change at 22°C (Figure S1B). We found a marginal *R*^2^ of 12%, with an additional 26% of the variation being explained by the random effects (conditional *R*^2^ = 38%).

When we separated unisexuals by their triploid biotype (LJJ, LLJ), we found that *V*CO_2_ increased with mass and temperature (Figure 2A). We also found that biotypes had different metabolic rates across temperatures, with unisexual biotypes exhibiting metabolic rates most similar to the sexual species with which they share the greater number of chromosome sets, specifically at the extreme temperatures (e.g. LJJ similar to *A. jeffersonianum* (JJ) but not to LLJ; Figure 2A; Table S1). For the Δ*V*CO_2_ analysis, Δ*V*CO_2_ was positively related to mass and was greatest at 22°C. We also found a significant effect of the interaction between temperature and biotype on Δ*V*CO_2_, with Δ*V*CO_2_ varying across biotypes and temperatures (Figure 2B). Follow-up pairwise analyses revealed a difference between JJ and LLJ at 6 for the acclimation analysis (*p* = 0.03), but there were no other pairwise differences between biotype pairs (*p* > 0.05; Table S1). Using AIC model comparison, we also found that models with biotype were more supported by the data than models that used species for the trait analysis (species: AIC = 281.18; biotype: AIC = 225.78) and acclimation analysis (species: AIC = 2922.08; biotype: AIC = 2901.88).

**Figure 2.**
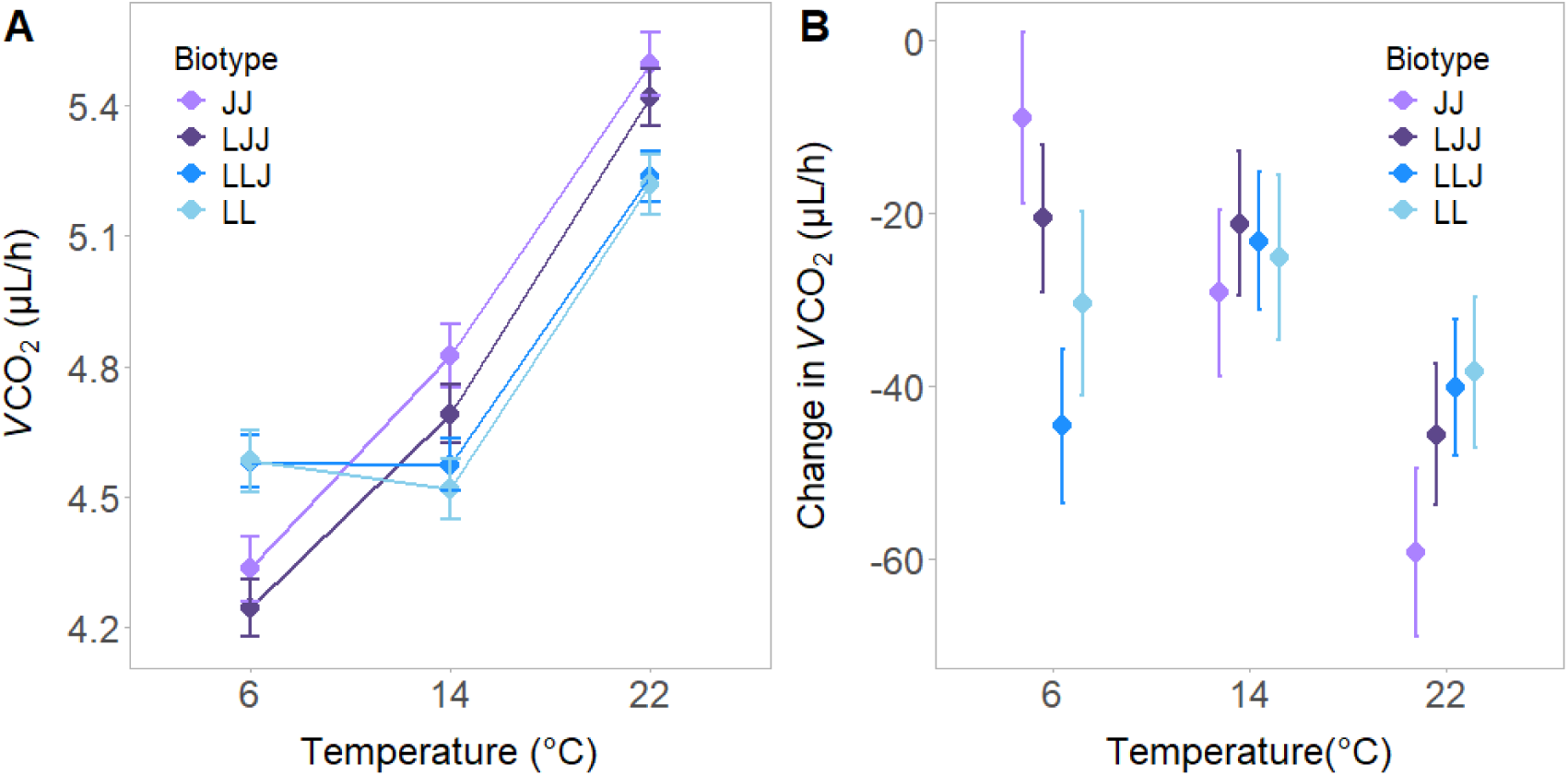
Variation in metabolic rate (*V*CO_2_) across temperature and biotype. (A) The effects of temperature on log-scaled *V*CO_2_ separated by biotype. (B) The effects of temperature on the change in *V*CO_2_ separated by biotype. Negative values in acclimation plot (B) show that salamanders acclimated by lowering their *V*CO_2_ following the acclimation treatment. Data for all panels are shown as the adjusted mean ± SEM, incorporating the effects of all variables in the model.

### Total resistance to water loss (r_T_)

For total resistance to water loss (*r_T_*), we found larger individuals and warmer temperatures had a higher *r_T_* (Figure S2A). Additionally, the interaction between temperature and species affected *r_T_*, with *A. laterale* having the lowest thermal sensitivity of resistance (Figure S2A). We also found that our treatments and mass explained 97% of the variation in our data. When analyzing Δ*r_T_*, we found that total resistance to water loss declined by 3% following exposure to the 20□- acclimation chamber (Figure S2B). We also found that Δ*r_T_* was greater in *A. jeffersonianum* than *A. laterale* or unisexuals (Figure S2B). Mass and treatments accounted for 7% of the variation, as shown by the marginal *R*^2^ value, with an additional 10% being explained by the random effects (conditional *R*^2^ = 17%).

When we separated unisexuals by their biotype, we found that *r_T_* was positively related to body mass, with larger individuals having the highest resistance, and *r_T_* increased with temperature (Figure 3A). The interaction between biotype and temperature also influenced *r_T_*, with LJJ and *A. jeffersonianum* (JJ) having the highest thermal sensitivity of resistance (Figure 3A). Similar to the *V*CO_2_ results, unisexual species exhibited a *r_T_* more similar to the sexual species they are more genetically like, specifically at the two extreme temperatures (Figure 3A; Table S1). For the Δ*r_T_* analysis, there was no effect of the interaction between temperature and biotype. However, there were differences among biotypes, with *Ambystoma jeffersonianum* (JJ) having the highest Δ*r_T_* value (Figure 3B). Using AIC model comparison, we found that models with biotype were more supported by the data than models that used species for the trait analysis (species: AIC = 2320.77; biotype: AIC = 2208.51). However, both models had similar AIC values for the change in trait analysis (species: AIC = 1185.87; biotype: AIC = 1184.54).

**Figure 3.**
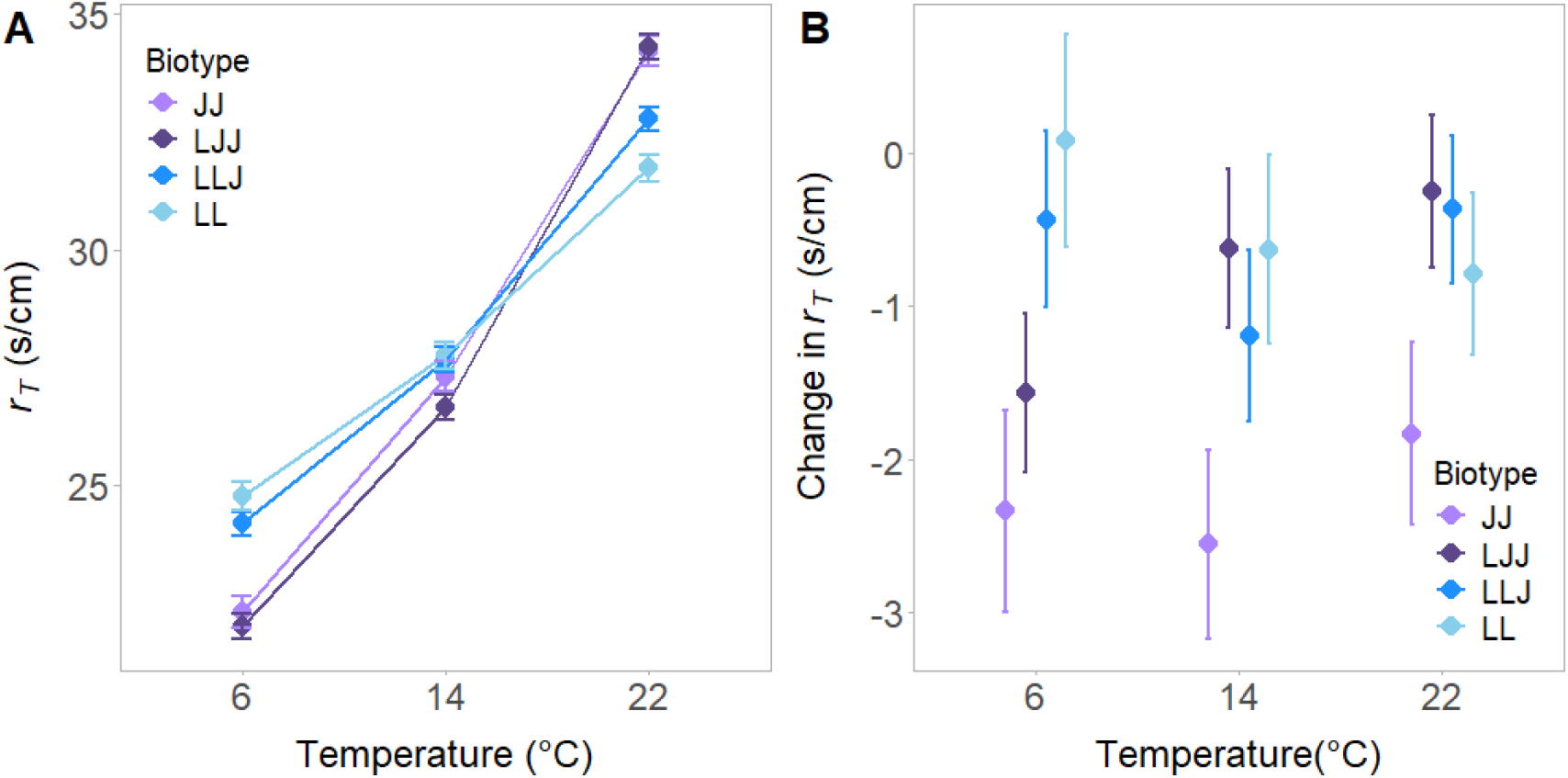
Variation in total resistance to water loss (*r_T_*) across temperature and biotype. (A) The effects of temperature on *r_T_* separated by biotype. (B) The effects of temperature on the change in *r_T_* separated by biotype. Negative values in the acclimation plot (B) show that salamanders acclimated by lowering their *r_T_* following the acclimation treatment. Data for all panels are shown as the adjusted mean ± SEM, incorporating the effects of all variables in the model.

### Respiration efficiency (WGER)

Respiration efficiency and mass were positively related, as were efficiency and temperature (Figure S3A). Furthermore, *A. jeffersonianum* had the highest respiration efficiency. We also found that the interaction between temperature and species influenced efficiency, with *A. laterale* having the least variation in efficiency across temperatures (Figure S3A). Our treatments and mass accounted for 83% of the variation, based on the marginal *R*^2^. In the ΔWGER analysis, we found that salamanders were 18% less efficient following the acclimation period, and they had the largest decrease in efficiency at 22°C (Figure S3B). Additionally, the interaction between temperature and species affected efficiency, with *Ambystoma jeffersonianum* having the greatest fluctuation in ΔWGER across temperatures (Figure S3B). Lastly, we found a marginal *R*^2^ of 15%, with an additional 25% of the variation being explained by the random effects (conditional *R*^2^ = 40%).

When we separated the unisexual group by biotype, we found that larger individuals had the highest respiration efficiency, and respiration efficiency increased as temperatures warmed (Figure 4A). We also found that the interaction between temperature and biotype was significant. Specifically, *A. jeffersonianum* and LJJ exhibited similar efficiencies at 6□ and 22□, as did *A. laterale* and LLJ (Figure 4A; Table S1). For ΔWGER, we found that all values were negative, indicating that individuals had lower respiratory efficiency following the acclimation treatment (Figure 4B). Additionally, ΔWGER was positively related to both mass and temperature. We also found an effect of the interaction between biotype and temperature on ΔWGER, with acclimation capacity varying by biotype and temperature (Figure 4B). However, the pairwise comparison did not identify any significant differences between biotypes in the ΔWGER analysis (Table S1). Using AIC model comparison, we found that models with biotype were more supported by the data than models that used species for the trait analysis (species: AIC = 217.35; biotype: AIC = 158.96); however, models with species were more supported by the data than models that used biotype for the change in trait analysis (species: AIC = −2667.24; biotype: AIC = −2623.78).

**Figure 4.**
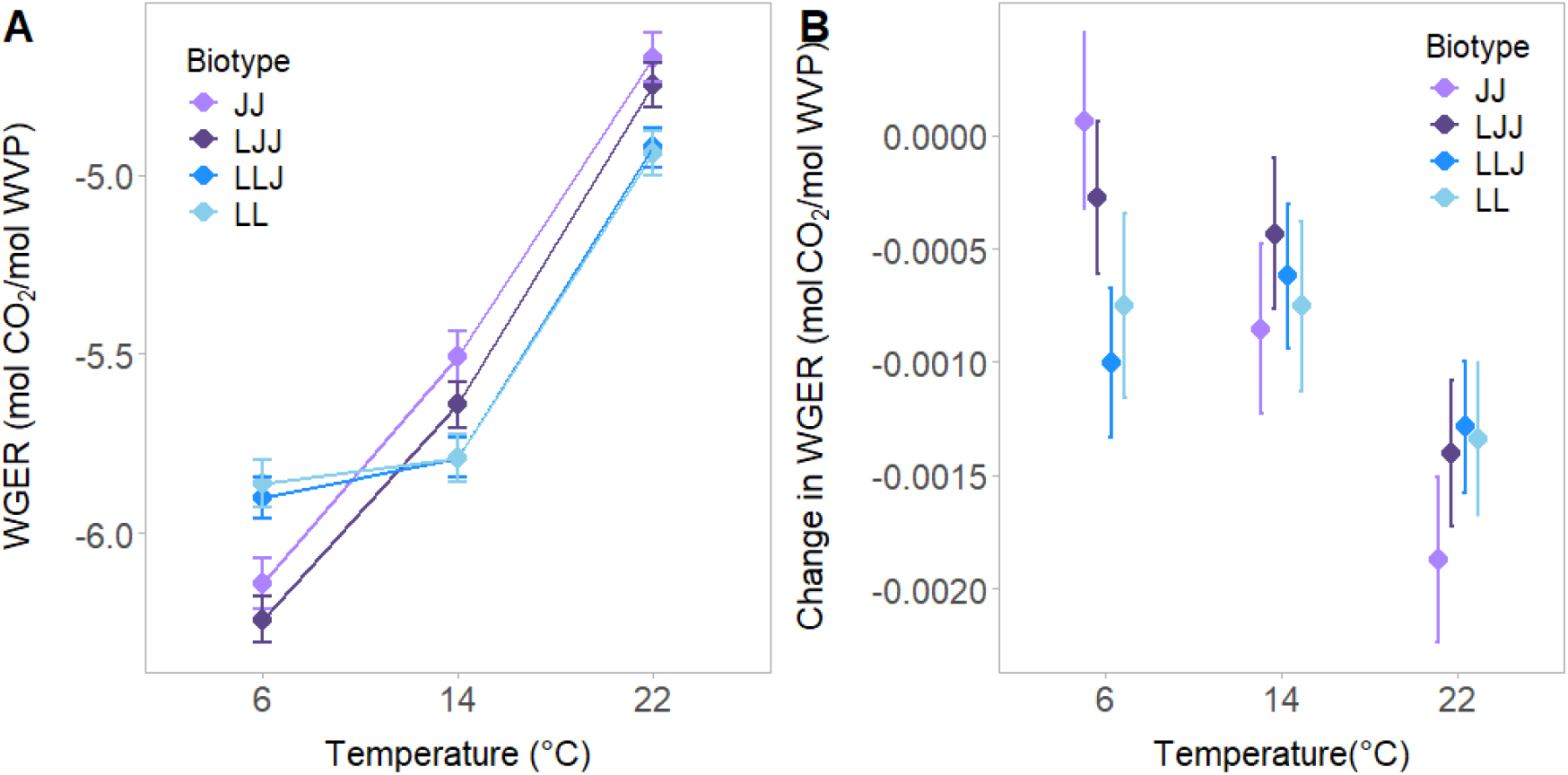
Variation in respiration efficiency (WGER) across temperature and biotype. (A) The effects of temperature on log-scaled WGER separated by biotype. (B) The effects of temperature on the change in WGER separated by biotype. Negative values in acclimation plot (B) show that salamanders acclimated by lowering their WGER following the acclimation treatment. Data for all panels are shown as the adjusted mean ± SEM, incorporating the effects of all variables in the model.

### Trade-offs between gas exchange and water loss

We also found a trade-off between the change in *r_T_* and *V*CO_2_, in which the change in resistance was negatively related to the change in *V*CO_2_ (Figure 5). We did not find any differences in the strength of the trade-off between species (Figure 5A) or temperature (Figure 5B). When separating unisexuals by biotype, we also found a significant trade-off between the change in *r_T_* and *V*CO_2_. However, we did not find any variation in the strength of the trade-off by biotype (Figure 5C). Using AIC model comparison, we found that models with biotype were more supported by the data than models that used species for the tradeoff analysis (species: AIC = 2536.86; biotype: AIC = 2502.35).

**Figure 5.**
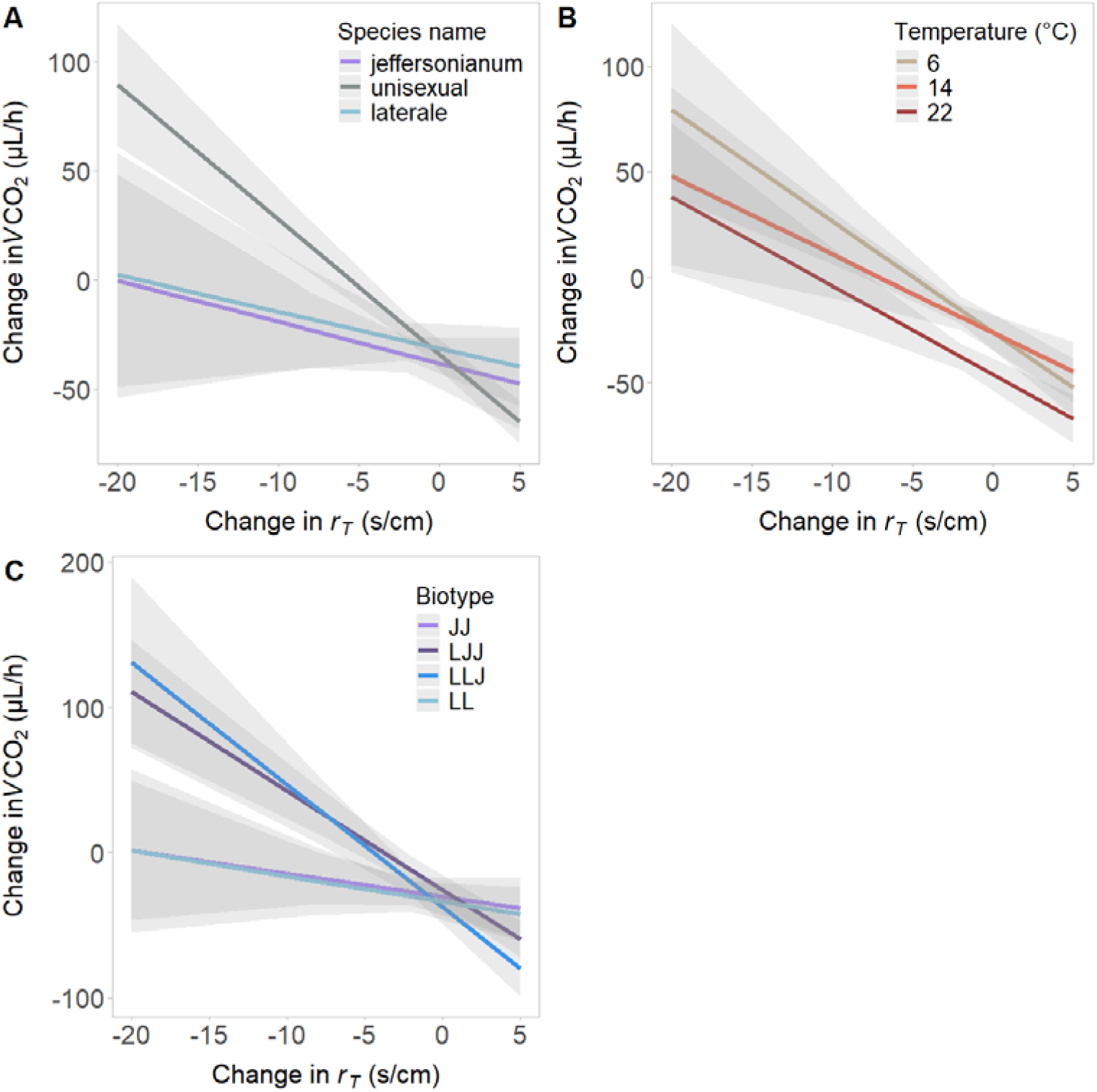
Trade-off between the acclimation of metabolic rate (*V*CO_2_) and change in total resistance to water loss (*r_T_*) across species (A), temperature (B), and biotype (C). Data for all panels are shown as adjusted mean ± SEM, incorporating the effects of all variables in the model.

## Discussion

The unisexual salamander system combines two potential consequences of allopolyploidization that may negatively impact metabolic processes: genetic incompatibilities from the interactions between divergent genomes and larger cell size via polyploidy. Our initial predictions were that unisexuals, regardless of biotype, would have reduced metabolic rate and respiration efficiency due to mitonuclear mismatch and lower metabolic and water loss rates due to cell size. However, we found that physiological performance of unisexual polyploids mirrored the most representative species in their sub-genome composition (Figure 1A, 2A, 3A), and unisexuals exhibited similar acclimation capacity compared to the sexual species (Figure 1B, 2B, 3B). Therefore, we found no evidence of allopolyploidization influencing physiological performance in unisexual salamanders, similar to other studies investigating metabolic and water loss rates in this system (Licht and Bogart, 1990; Licht and Lowcock, 1991) but unlike studies that investigated endurance (Denton et al., 2017). To the best of our knowledge, this study is the first to include both sexual species that contribute genomes to the unisexual lineage (*A. laterale* and *A. jeffersonianum*) and subsequently the first to show that subgenome composition may play a larger role than ploidy in estimating physiological performance in this system. These results highlight the importance of including the performance of the parental species when measuring and interpreting the performance of hybrid offspring.

Accounting for the subgenome composition in hybrids may be necessary for understanding variation in hybrid performance across traits. Unisexual *Ambystoma* have greater performance in certain traits (i.e., tissue regeneration) compared to congenerics (Saccucci et al. 2016), but they have lower performance in traits related to endurance compared to sexual species, possibly due to mitonuclear mismatch (Denton et al. 2017). Alternatively, the unisexual *Ambystoma* salamanders in this study broadly display intermediate values between the sexual species and mirror the values of their major subgenome when comparing unique biotypes (i.e. LLJ is more like *A. laterale* than LJJ). The inconsistencies of unisexual performance compared to sexual congenerics across these studies may be due to the differences in the genomic array of the unisexuals in each study. While the unisexual lineage is very old (Bogart et al. 2007; Bi and Bogart 2010), it readily adds genomic variation from nearby species into its genome (Bi et al. 2008; Gibbs and Denton 2016; Denton et al. 2018). Therefore, variation in unisexual performance across different traits (e.g., metabolism and total resistance to water loss, this study; tissue regeneration, Saccucci et al. 2016; and endurance/dispersal rates, Denton et al. 2017) likely depends on the genomic composition of each individual and the relative performance of the sexual species from which each genome was received. Additionally, our study measured resting metabolic rate as opposed to active or maximal metabolic rate (as measured in Denton et al. 2017). Metabolic rate during endurance trials relies on a higher rate of ATP production to fuel the greater energetic demand (Clarke and Fraser, 2004); therefore, mitonuclear mismatch may have more of an impact during activity since it reduces the production of ATP. Therefore, accounting for the genomic array of the unisexuals and the physiological traits being measured is necessary to understand hybrid performance across studies.

The similarities between unisexuals and sexual species in this study provide support for the persistence of unisexuals across millennia. Unisexuals commonly underperform sexual congenerics, having lower fecundity (Uzzell 1969), being avoided by sexual males in mate-choice trials (Dawley and Dawley, 1986), and having lower endurance and dispersal rates (Denton et al., 2017) compared to sexual species. Unisexuals also require the presence of sympatric sexual species to reproduce, with unisexual populations declining when sexual species are absent (Bogart et al., 2016). In fact, the persistence of unisexuals may even rely on cyclical patterns of population fluctuations between competing unisexuals and sexual species (Greenwald et al., 2016). Thus, these combined effects suggest that unisexuals should be outcompeted by sexual species and not be able to persist. However, unisexuals exhibit faster growth rates (Licht and Bogart, 1987), have a competitive advantage over certain congenerics as larvae (outcompete *A. laterale* larvae but not *A. texanum*; Brodman and Krouse, 2007), can potentially drive sexual populations to extinction (Clanton, 1934), and as shown here, do not exhibit any differences in metabolic rate, water loss rate, or acclimation capacity. Understanding the coexistence between sexual and unisexual species may require linking physiology to organismal performance across ecological and environmental contexts, especially given that the species in this study are considered threatened or protected in many parts of their range (Canada’s Species at Risk Act, 2002; Illinois DNR, 2020; Ohio DNR, 2022;).

Species responses to environmental change may be partly explained by the environmental conditions experienced throughout their geographic range (Navas, 1996; Wilson, 2001; Lancaster et al., 2020). *Ambystoma laterale* is the most cold-tolerant salamander in North America (Storey and Storey 1992) and is primarily found in higher latitudes and previously glaciated regions (Figure 1B), so higher performance at cooler temperatures may be an adaptive advantage for this species. We observed higher metabolic rates, resistance to water loss, and respiration efficiency in *A. laterale* at cool temperatures, which might facilitate prolonged activity while minimizing desiccation risk at temperatures they experience more regularly. *Ambystoma laterale* also have the most widespread geographic range and therefore experience a broader thermal gradient (Figure 1B). Their lower thermal sensitivity to acute changes in temperature (Figures 2-4) may lessen any negative effects of higher thermal variation by minimizing energetic costs at different temperatures. Alternatively, *A. jeffersonianum* is subject to warmer average temperatures, which could explain the higher metabolic rate, resistance to water loss, and respiration efficiency at higher temperatures. An increase in these traits at warmer temperatures may facilitate a greater capacity for work while minimizing desiccation risk. *Ambystoma jeffersonianum* also have a much smaller geographic range (Figure 1B), providing a potential explanation for their relatively higher thermal sensitivity of physiological traits. Unisexuals experience a transitional climate, overlapping both *A. laterale* and *A. jeffersonianum* ranges; therefore, their intermediate phenotypes may be the result of evolving in this geographic region (Figure 1B) or simply the additive effects of their sub-genomes (McElroy et al. 2017). Geographic locations may also explain unisexual performance, since LLJ biotypes are often found in *A. laterale* populations, and LJJ are found in *A. jeffersonianum* populations (Figure 1B; Appendix 1). These interspecific differences and the environmental conditions that explain them could inform predictions on how environmental change will affect species physiological responses, performance, and fitness (Riddell et al., 2018).

Including phenotypic measurements of parental species is imperative for an accurate interpretation of hybrid performance. In this study, we discovered that subgenome composition was highly indicative of hybrid trait response, with biotypes performing more like the parental species of their major subgenomes. Although we found no effect of allopolyploidization on the traits studied here, measuring other metrics of performance, such as maximal metabolic rate, could better elucidate the effects of allopolyploidization on hybrid performance. Therefore, further investigation into phenotypic variation and the mechanistic drivers of this variation could give insight into not only the persistence of certain lineages throughout millennia, but also the persistence and vulnerability of various species towards current and future environmental change.

