## Appendix1_supplement for "Genome composition predicts physiological responses to temperature in polyploid salamanders"

**Appendix 1: Effects of collection site on physiological response within biotypes**

*Statistical analysis*

Because biotype and collection site are confounded (specific biotypes are only found in specific locations; Fig. 1), we could not account for site effects in our global model. Therefore, we ran additional analyses to determine the effects of collection site on performance within each biotype. We subset our master dataset into four different datasets, one for each biotype (LL, LLJ, LJJ, JJ). We then ran the following analyses for each separate biotype. For metabolic rate, resistance to water loss and respiration efficiency (WGER), we treated the physiological trait as the response variable, mass as a covariate, and temperature treatment, time period (before and after acclimation), and collection site as factors. We also incorporated a temperature by time period interaction, similar to our global model. We log-transformed metabolic rate, respiration efficiency, and mass for these models to meet assumptions of normalcy. For the tradeoff analysis, we included collection site as a factor to determine the effect of site on the trade-off between the change in resistance to water loss and the change in metabolic rate. All models included mass as a covariate. Salamander ID nested inside batch number was included as a random effect for all models except the trade-off analysis for JJ because it did not account for any additional variation in the model. Year collected was not included as a random effect in any model because it did not account for any additional variation. Whenever county had a significant effect on the results, we followed up the analyses with pairwise comparisons using the emmeans package.

*Results*

We found that collection site had an effect on metabolic rate for each biotype (LL: F_3_ = 7.61, *p* = 0.0005, *ω^2^* = 0.15; LLJ: F_3_ = 3.14, *p* = 0.0005, *ω^2^* = 0.34; LJJ: F_3_ = 4.61, *p* = 0.009, *ω^2^* = 0.26; JJ: F_2_ = 3.75, *p* = 0.04, *ω^2^* = 0.23). Specifically, LL in Livingston, MI had a significantly lower metabolic rate than those in Marquette, MI (*p* = 0.02), and LLJ in Livingston, MI had a significantly lower metabolic rate than those in Addison, VT (*p* = 0.008) and Calumet, WI (*p* = 0.0008). Therefore, for LLJ and LL biotypes, more southern populations (Livingston, MI) had lower metabolic rates than northern populations.

For LJJ, salamanders collected in Addison, VT had a significantly lower metabolic rate than those collected from Delaware, OH (*p* = 0.03) and Litchfield, CT (*p* = 0.01), and salamanders collected from Campbell, KY had a lower rate than those from Litchfield, CT (*p* = 0.04). JJ salamanders from Campbell, KY also had a lower metabolic rate than salamanders from Litchfield, CT (*p* = 0.03). Therefore, for LJJ, the northern-most population had the lowest metabolic rate (Addison, VT); however, the second lowest metabolic rate for LJJ was from the southern-most population, and JJ salamanders collected from this population (Campbell, KY) also had the lowest metabolic rate. When combined with results from LL/LLJ populations, we found that more southern populations typically exhibit lower metabolic rates than northern populations, except for LJJ salamanders collected from Addison, VT (Figure A1).

We found that collection site had no effect on resistance to water loss (LL: F_3_ = 1.29, *p* = 0.29, *ω^2^* = 0.03; LLJ: F_3_ = 0.70, *p* = 0.56, *ω^2^* < 0.001; LJJ: F_3_ = 0.45, *p* = 0.72, *ω^2^* < 0.001; JJ: F_2_ = 0.56, *p* = 0.58, *ω^2^* < 0.001) (Figure A2).

For respiration efficiency, collection site had an effect on performance for LLJ (F_3_ = 7.14, *p* = 0.0008, *ω^2^* = 0.34) and LJJ (F_3_ = 4.18, *p* = 0.01, *ω^2^* = 0.24), but not for LL (F_3_ = 2.90, *p* = 0.05, *ω^2^* = 0.14) and JJ (F_2_ = 2.96, *p* = 0.08, *ω^2^* = 0.18). Similar to the metabolic rate outputs, LLJ salamanders collected from Livingston, MI had a lower respiration efficiency than those collected from Addison, VT (*p* = 0.01) and Calumet, WI (*p =* 0.001). LJJ salamanders from Addison, VT had a lower respiration efficiency than those from Delaware, OH (*p* = 0.04) and Litchfield, CT (*p* = 0.02). Therefore, southern populations of LLJ exhibited the lowest efficiency, but the northernmost population of LJJ had the lowest efficiency (Figure A3)

The effect of collection site on the trade-off also varied by biotype, having a significant effect for LJJ (F_3_ = 5.39, *p* = 0.002, *ω^2^* = 0.16) but no effect for LLJ (F_3_ = 1.30, *p* = 0.28, *ω^2^* = 0.02), LL (F_3_ = 1.81, *p* = 0.15, *ω^2^* = 0.04), or JJ (F_3_ = 1.55, *p* = 0.23, *ω^2^* = 0.03). LJJ individuals from Delaware, OH exhibited the strongest trade-off, while those collected from Campbell, KY and Addison, VT have little to no evidence of a trade-off. Thus, LJJ individuals with lower metabolic rates (those collected from Campbell, KY and Addison, VT) exhibited less evidence of a trade-off, indicating that they may be able to decouple metabolic rate and water loss rates, unlike salamanders with higher metabolic rates (Delaware, OH; Figure A4).

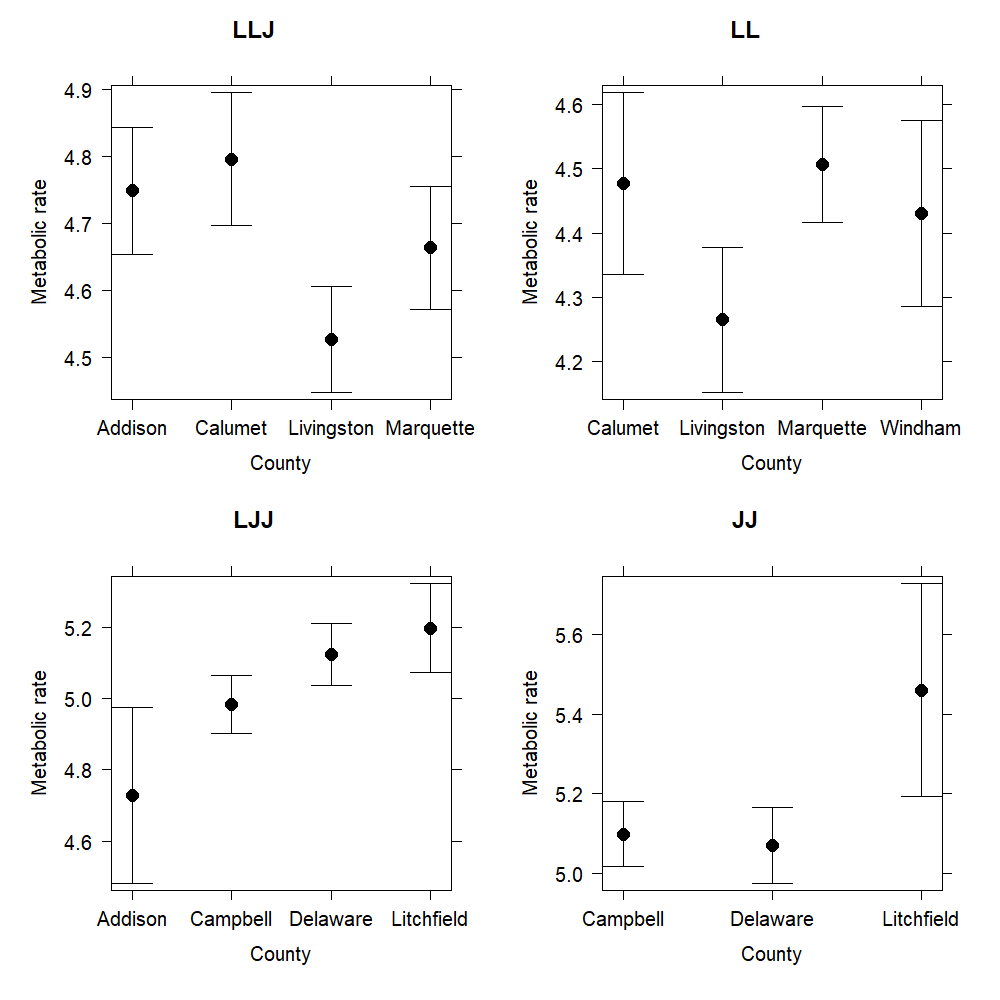

Figure A1: Log-scaled metabolic rates of unisexual (LLJ and LJJ) and sexual (LL and JJ) biotypes across collection sites. LLJ and LL salamander from Livingston, MI had a lower average metabolic rate than the other collection sites. LJJ individuals from Addison, VT exhibited the lowest metabolic rate, and LJJ and JJ salamanders from Campbell, KY had lower metabolic rates than those from Litchfield, CT. Data is shown as the adjusted mean ± 95% CI.

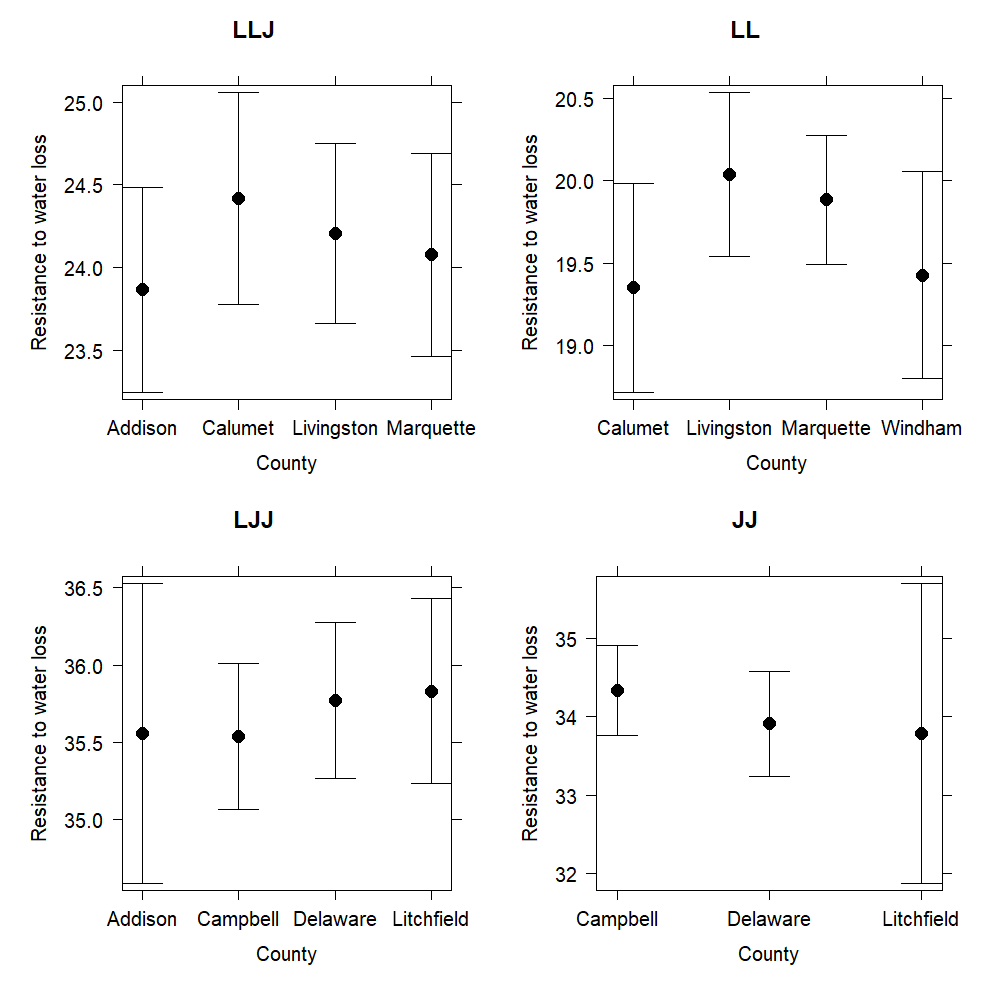

Figure A2: Resistance to water loss rates of unisexual (LLJ and LJJ) and sexual (LL and JJ) biotypes across collection sites. There was no significant differences in resistance to water loss across collection sites. Data is shown as the adjusted mean ± 95% CI.

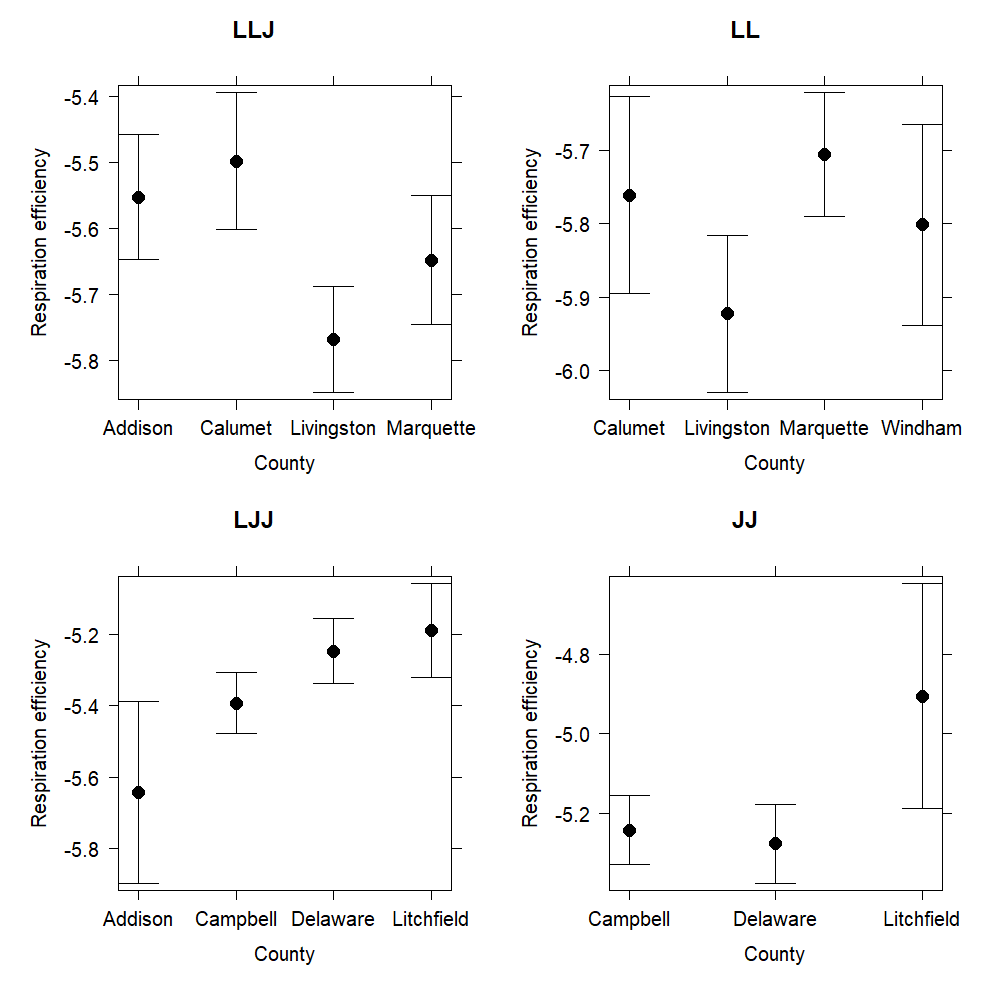

Figure A3: Log-scaled respiration efficiency of unisexual (LLJ and LJJ) and sexual (LL and JJ) biotypes across collection sites. LLJ salamanders from Livingston, MI and LJJ salamanders from Addison, VT had the lowest average respiration efficiencies. There was no effect of county on respiration efficiency for LL and JJ individuals. Data is shown as the adjusted mean ± 95% CI.

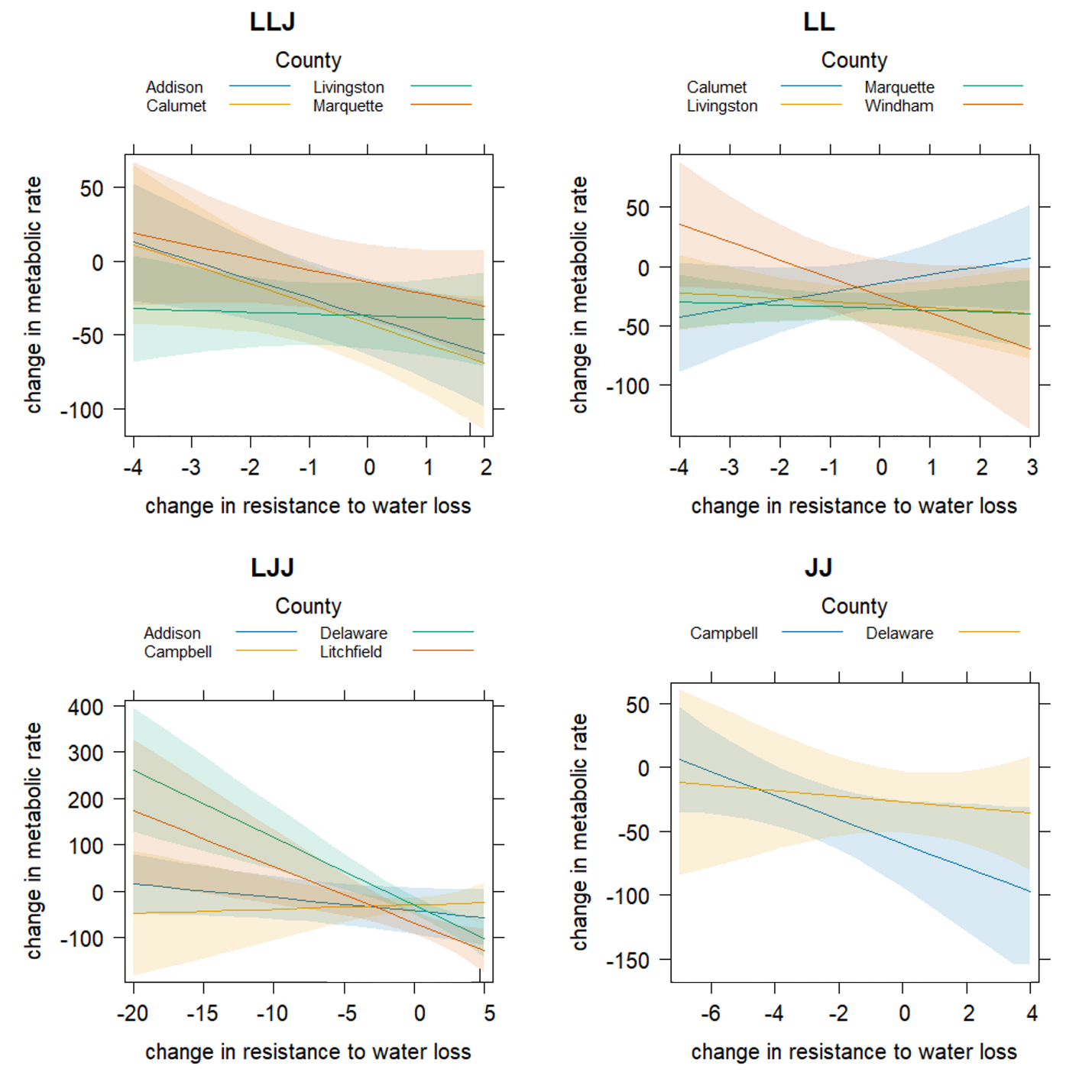

Figure A4: Trade-off between the change in resistance to water loss and the change in metabolic rate across unisexual biotypes (LLJ and LJJ) and sexual biotypes (LL and JJ) across collection sites. The trade-off in LJJ individuals was significantly different by county, with Delaware, OH having the strongest trade-off and Campbell, KY and Addison, VT exhibiting almost no trade-off. There was no effect of county on the trade-off for LLJ, LL, and JJ populations. Data is shown as the adjusted mean ± 95% CI.
