## Supplemental_tables_and_figures for "Genome composition predicts physiological responses to temperature in polyploid salamanders"

**Supplemental table and figures**

Table S1 Results of pairwise comparisons between biotypes

| Trait | | Temperature (℃) | Comparison | *p*-value |
| --- | --- | --- | --- | --- |
| *V*CO_2_ | | 6 | JJ – LJJ | 0.40 |
|  |  | 6 | JJ – LLJ | 0.006 |
|  |  | 6 | JJ – LL | 0.01 |
|  |  | 6 | LJJ – LLJ | < 0.001 |
|  |  | 6 | LJJ – LL | < 0.001 |
|  |  | 6 | LLJ – LL | 1 |
|  |  | 14 | JJ – LJJ | 0.11 |
|  |  | 14 | JJ – LLJ | 0.002 |
|  |  | 14 | JJ – LL | 0.0003 |
|  |  | 14 | LJJ – LLJ | 0.26 |
|  |  | 14 | LJJ – LL | 0.06 |
|  |  | 14 | LLJ – LL | 0.77 |
|  |  | 22 | JJ – LJJ | 0.59 |
|  |  | 22 | JJ – LLJ | 0.001 |
|  |  | 22 | JJ – LL | 0.001 |
|  |  | 22 | LJJ – LLJ | 0.02 |
|  |  | 22 | LJJ – LL | 0.02 |
|  |  | 22 | LLJ – LL | 0.99 |
| Δ*V*CO_2_ | | 6 | JJ – LJJ | 0.69 |
|  |  | 6 | JJ – LLJ | 0.03 |
|  |  | 6 | JJ – LL | 0.46 |
|  |  | 6 | LJJ – LLJ | 0.20 |
|  |  | 6 | LJJ – LL | 0.90 |
|  |  | 6 | LLJ – LL | 0.61 |
|  |  | 14 | JJ – LJJ | 0.87 |
|  |  | 14 | JJ – LLJ | 0.96 |
|  |  | 14 | JJ – LL | 0.99 |
|  |  | 14 | LJJ – LLJ | 1 |
|  |  | 14 | LJJ – LL | 0.99 |
|  |  | 14 | LLJ – LL | 1 |
|  |  | 22 | JJ – LJJ | 0.56 |
|  |  | 22 | JJ – LLJ | 0.37 |
|  |  | 22 | JJ – LL | 0.37 |
|  |  | 22 | LJJ – LLJ | 0.96 |
|  |  | 22 | LJJ – LL | 0.93 |
|  |  | 22 | LLJ – LL | 1 |
| *r_T_* | 6 | | JJ – LJJ | 0.79 |
|  | 6 | | JJ – LLJ | < 0.001 |
|  | 6 | | JJ – LL | <0.0001 |
|  | 6 | | LJJ – LLJ | <0.0001 |
|  | 6 | | LJJ – LL | <0.0001 |
|  | 6 | | LLJ – LL | 0.23 |
|  | 14 | | JJ – LJJ | 0.25 |
|  | 14 | | JJ – LLJ | 0.78 |
|  | 14 | | JJ – LL | 0.70 |
|  | 14 | | LJJ – LLJ | 0.03 |
|  | 14 | | LJJ – LL | 0.04 |
|  | 14 | | LLJ – LL | 0.99 |
|  | 22 | | JJ – LJJ | 0.99 |
|  | 22 | | JJ – LLJ | 0.0006 |
|  | 22 | | JJ – LL | <0.0001 |
|  | 22 | | LJJ – LLJ | <0.0001 |
|  | 22 | | LJJ – LL | <0.0001 |
|  | 22 | | LLJ – LL | 0.001 |
| Δ*r_T_* | | | JJ – LJJ | 0.007 |
|  |  |  | JJ – LLJ | 0.03 |
|  |  |  | JJ – LL | 0.04 |
|  |  |  | LJJ – LLJ | 0.99 |
|  |  |  | LJJ – LL | 0.94 |
|  |  |  | LLJ – LL | 0.96 |
| WGER | 6 | | JJ – LJJ | 0.33 |
|  | 6 | | JJ – LLJ | 0.009 |
|  | 6 | | JJ – LL | 0.003 |
|  | 6 | | LJJ – LLJ | <0.0001 |
|  | 6 | | LJJ – LL | <0.0001 |
|  | 6 | | LLJ – LL | 0.93 |
|  | 14 | | JJ – LJJ | 0.10 |
|  | 14 | | JJ – LLJ | 0.0007 |
|  | 14 | | JJ – LL | 0.0012 |
|  | 14 | | LJJ – LLJ | 0.11 |
|  | 14 | | LJJ – LL | 0.15 |
|  | 14 | | LLJ – LL | 1 |
|  | 22 | | JJ – LJJ | 0.54 |
|  | 22 | | JJ – LLJ | 0.002 |
|  | 22 | | JJ – LL | 0.002 |
|  | 22 | | LJJ – LLJ | 0.03 |
|  | 22 | | LJJ – LL | 0.03 |
|  | 22 | | LLJ – LL | 0.99 |
| ΔWGER | 6 | | JJ – LJJ | 0.79 |
|  | 6 | | JJ – LLJ | 0.12 |
|  | 6 | | JJ – LL | 0.38 |
|  | 6 | | LJJ – LLJ | 0.36 |
|  | 6 | | LJJ – LL | 0.75 |
|  | 6 | | LLJ – LL | 0.94 |
|  | 14 | | JJ – LJJ | 0.66 |
|  | 14 | | JJ – LLJ | 0.96 |
|  | 14 | | JJ – LL | 0.99 |
|  | 14 | | LJJ – LLJ | 0.97 |
|  | 14 | | LJJ – LL | 0.89 |
|  | 14 | | LLJ – LL | 0.99 |
|  | 22 | | JJ – LJJ | 0.52 |
|  | 22 | | JJ – LLJ | 0.52 |
|  | 22 | | JJ – LL | 0.60 |
|  | 22 | | LJJ – LLJ | 0.99 |
|  | 22 | | LJJ – LL | 0.99 |
|  | 22 | | LLJ – LL | 0.99 |

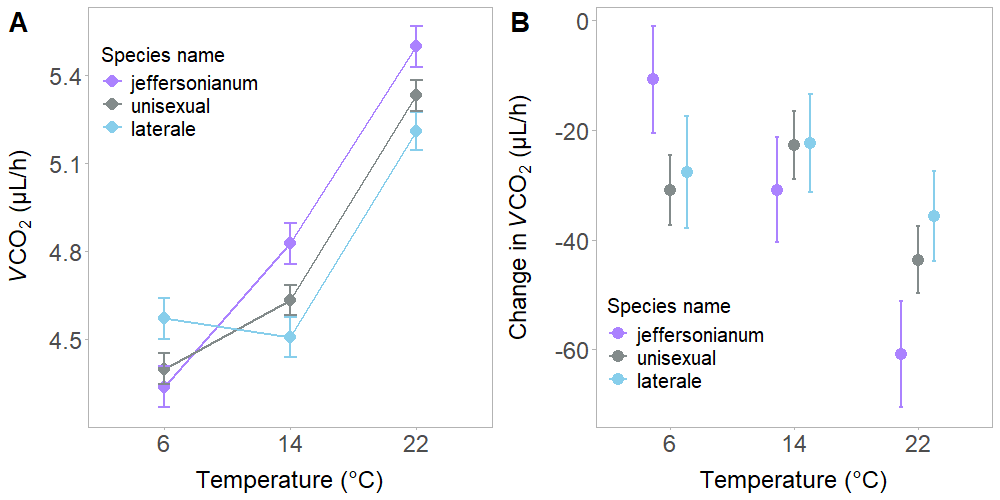

**Figure S1**. Variation in metabolic rate (*V*CO_2_) across temperature and species. (A) The effects of temperature on log-scaled *V*CO_2_ separated by species. (B) The effects of temperature on the change in *V*CO_2_ separated by species. Negative values in acclimation plot (B) show that salamanders acclimated by lowering their *V*CO_2_ following the acclimation treatment. Data for all panels are shown as the adjusted mean ± SEM, incorporating the effects of all variables in the model.

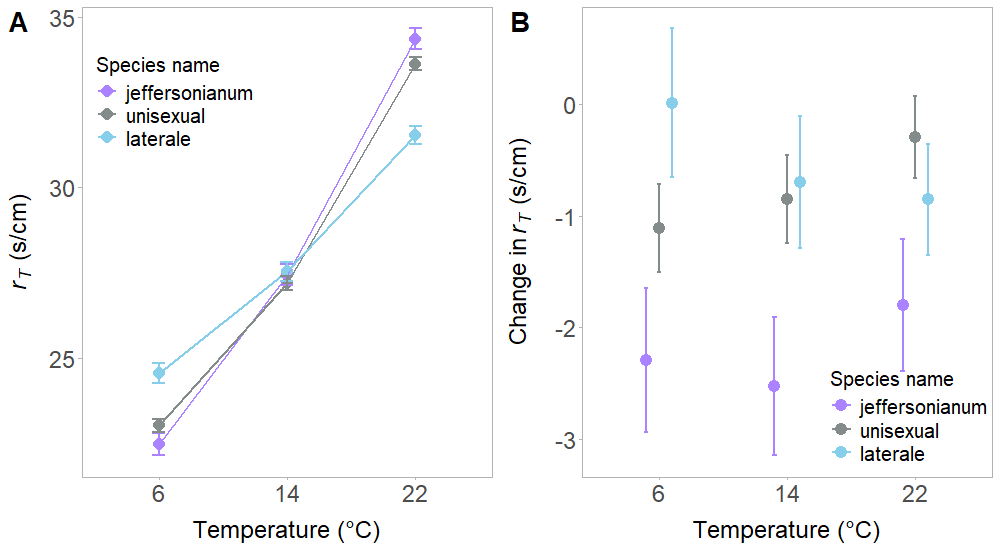

**Figure S2** Variation in total resistance to water loss (*r_T_*) across temperature and species. (A) The effects of temperature on *r_T_* separated by species. (B) The effects of temperature on the change in *r_T_* separated by species. Negative values in the acclimation plot (B) show that salamanders acclimated by lowering their *r_T_* following the acclimation treatment. Data for all panels are shown as the adjusted mean ± SEM, incorporating the effects of all variables in the model.

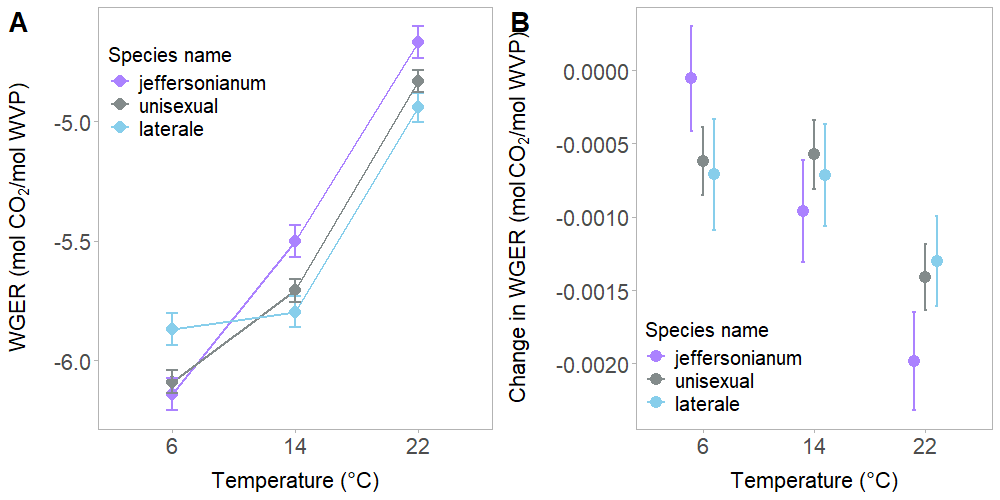

**Figure S3** Variation in respiration efficiency (WGER) across temperature and species. (A) The effects of temperature on WGER separated by species. (B) The effects of temperature on the change in WGER separated by species. Negative values in acclimation plot (B) show that salamanders acclimated by lowering their WGER following the acclimation treatment. Data for all panels are shown as the adjusted mean ± SEM, incorporating the effects of all variables in the model.
